# miRNAs Delivered by Prostate Cancer Extracellular Vesicles Coordinate the Regulation of Bone Metastasis

**DOI:** 10.64898/2026.01.23.700652

**Authors:** Lijuan Yu

**Author notes:** **Correspondence**: Lijuan Yu.

## Abstract

**Background:** We previously reported that extracellular vesicles derived from osteoblastic, osteoclastic, and mixed prostate cancer (PCa) cells promote osteoclast differentiation and inhibit osteoblast differentiation via the transfer of miR-92a-1-5p. However, at the same time, osteoblastic miRNAs also exist in PCa extracellular vesicles. In the present study, we focused on discovering (1) the roles of miR-375 and miR-148a-3p delivered by PCa extracellular vesicles in bone homeostasis and bone metastasis, and (2) an explanation for the co-existence of osteoblastic miRNAs and osteoclastic miRNAs in prostate cancer extracellular vesicles.

**Methods:** Conditional medium, miRNA mimics, and miRNAs overexpressed lentiviruses were employed to discover the roles of miR-375 and miR-148a-3p delivered by PCa extracellular vesicles in bone homeostasis and tumor growth. Target gene prediction and siRNAs were employed to discover the target gene(s) for miR-375 and miR-148a-3p. Droplet digital PCR (ddPCR) of the PCa extracellular vesicle miRNAs were utilized to evaluate miRNA expression at different metastatic phases.

**Results:** Conditional medium from prostate cancer cell culture promoted osteoblast differentiation, as confirmed by ALP staining and Alizarin red staining. miR-375 and miR-148a-3p promoted osteoblast differentiation *in vitro* by reducing *KLF4* expression, associated with increased osteoblast function as shown by ALP staining, Alizarin red staining, and *Alp* mRNA expression. *In vivo*, miR-375 and miR-148a-3p overexpressing MDA PCa 2b cells promoted osteoblastogenesis and tumor growth, which were confirmed by micro computed tomography and *in vivo* imaging. siRNA targeting *KLF4* similarly enhanced osteoblast function and tumor cell proliferation. Based on the miRNA ddPCR data for PCa extracellular vesicles, osteoclastic and osteoblastic miRNAs existed in different bone metastatic phases.

**Conclusions:** These findings suggest that miR-375 and miR-148a-3p delivered by PCa extracellular vesicles regulate osteoblast function and tumor growth via targeting *KLF4*. Osteoclastic miR-92a-1-5p is active during the early phases of bone metastasis, while osteoblastic miR-375 and miR-148a-3p function during the late phases.

## Background

Bone metastasis represents the primary cause of mortality among patients with advanced prostate cancer (PCa) [1]. As the densest tissue within the human body, bone constitutes the most prevalent site for PCa metastasis [2]. In accordance with the “seed and soil” hypothesis, both PCa cells and bone tissue actively contribute to the bone metastatic microenvironment [3, 4]. PCa cells release cytokines and extracellular vesicles (EVs) to modulate the bone environment, while PCa cells residing in the bone secrete growth factors that facilitate tumor proliferation, thereby establishing a “vicious cycle” [5-7]. The homeostatic balance between osteoblasts and osteoclasts plays a critical role in regulating bone metastasis.

Recent research has elucidated the significant role of EVs and their associated cargoes in mediating intercellular and interorgan communication [8-11]. In the context of bone-metastatic PCa, numerous studies have identified miRNAs delivered by PCa cells as contributors to bone metastasis. In 2017, our research group demonstrated that miR-141-3p, conveyed via MDA PCa 2b derived exosomes, facilitates the osteoblastic differentiation of human osteoblasts and promotes tumor-induced osteoblastic bone metastasis [12]. In 2018, Hashimoto et al. subsequently reported that miR-940, also delivered by PCa cells, enhances the osteogenic differentiation of human mesenchymal stem cells *in vitro* and induces osteoblastic lesions *in vivo* [13]. In 2021, we further reported that miR-92a-1-5p, which is upregulated 30-fold in MDA PCa 2b derived EVs compared to EVs derived from normal prostate cells, augments osteoclastic activity in Raw264.7 cells and bone marrow macrophages, while inhibiting the osteoblastic differentiation of MC3T3-1 cells [14]. Clinically, PCa bone metastasis is mainly osteoblastic, but the coexistence of osteoblastic and osteoclastic miRNAs in PCa EVs remains unexplained.

In our previous work, we demonstrated that EVs derived from osteoblastic (MDA PCa 2b), osteoclastic (PC3), and mixed (C4-2) PCa cells facilitate osteoclast differentiation while inhibiting osteoblast differentiation [14]. Notably, the microRNA miR-92a-1-5p was identified as a significant factor that enhances osteoclast differentiation and suppresses osteoblastogenesis. Additionally, we reported that PCa EVs engineered to carry miR-92a-1-5p modulate osteoclast function through the MAPK1/FOXO1 signaling pathway [15]. Furthermore, we identified two osteoblastic microRNAs, miR-375 and miR-148a-3p, which are conveyed by PCa EVs. The present study aims to (1) elucidate the functional roles and mechanisms of action of miR-375 and miR-148a-3p delivered by PCa EVs in bone homeostasis and metastasis, and (2) further explore the rationale behind the coexistence of osteoblastic and osteoclastic microRNAs in the bone metastatic process.

## Material and Methods

### Cell culture

The human PCa cell line (MDA PCa 2b) was purchased from American Type Culture Collection (ATCC, USA) and cultured in F12K medium (ATCC) supplemented with 20% fetal bovine serum (FBS), according to the manufacturer’s instructions. The MC3T3-E1 clone 4 cell line (ATCC) was maintained in α-MEM (Invitrogen, USA) containing 10% FBS. All culture media were replaced with fresh medium every 3 days and supplemented with 100 U/ml penicillin and 100 μg/ml streptomycin (HyClone, USA).

### Conditioned medium preparation

Conditioned medium was prepared as previously described [16]. Briefly, 5 × 10^6^ MDA PCa 2b cells were cultured in 75-cm^2^ flasks with F12K medium and 20% FBS for 12 hours. The medium was then switched to F12K with 0.5% FBS, with or without 20 μM GW4869. After 48 hours, the supernatant was collected, centrifuged at 2,500 rpm for 10 minutes to remove debris, and filtered through a 0.2-μm filter before experimental use.

### Lentivirus transfection

Lentiviral plasmids encoding miR-375, miR-148a-3p, and a negative control were synthesized by Genepharma (Lot: L2021-SH3, Shanghai, China). The MDA PCa 2b and MC3T3-E1 cell lines were subsequently transfected with the lentivirus (LV-U6-shRNA-CMV-Luciferase17-Puro) at a multiplicity of infection of 10. Following transfection, the cells were subjected to selection with 4 μg/ml puromycin (Solarbio, China) for six days.

### *In vitro* osteogenic induction of MC3T3-E1 cells

MC3T3-E1 cells were initially seeded into 24-well plates at a density of 1 × 10^4^ cells per well on day 0 and cultured in α-MEM until reaching 80-90% confluence. Subsequently, the medium was switched to an osteogenic differentiation medium (Cyagen Biosciences, China). Following a 21-day induction period, Alizarin Red S staining was conducted as previously described [14]. Specifically, the cells were fixed with 4% paraformaldehyde for 15 minutes, incubated with Alizarin Red S (Cyagen Biosciences) for 3 minutes, and subsequently rinsed with deionized water on day 21. To quantify mineralization, a 2% solution of hexadecyl pyridinium chloride monohydrate was applied, and absorbance was measured at 490 nm using a plate reader. For alkaline phosphatase (ALP) staining, the cells were stained 7 days post-osteogenic induction utilizing the BCIP/NBT alkaline phosphatase chromogenic kit (Beyotime) in accordance with the manufacturer’s instructions.

### Cell proliferation assay

A cell proliferation assay was conducted utilizing the Cell Counting Kit 8 (CCK8, Dojindo, Japan), following the methodology previously outlined [17]. Specifically, cells were seeded into 96-well plates at a density of 1 × 10^3^ cells per well on day 0. Cell density was assessed at various time points (0, 24, 48, 72, and 96 hours) by introducing 10 μL of CCK-8 reagent into the cell culture medium and incubating at 37°C for 1 hour. Subsequently, the optical density at 450 nm was measured using a microplate reader.

### Quantitative real-time PCR (qPCR)

To investigate miRNA expression within EVs, RNA was extracted utilizing the Total Exosome RNA and Protein Isolation Kit (Invitrogen). Subsequently, the Mir-X™ miRNA First Strand Synthesis Kit (Takara, Japan) and SYBR® Premix Ex Taq™ II (Takara) were employed for quantitative PCR (qPCR) to measure the expression levels of miR-375 and miR-148a-3p. Cel-miR-39 (Qiagen, German) served as a spike-in control. The miRNA qPCR primer sets were procured from RiboBio (Beijing, China).

### Luciferase reporter assay

Cells were co-transfected with a luciferase vector harboring either the wild-type or mutant 3D-untranslated region (3D-UTR) of *KLF4*, alongside miRNA mimics or a miR-Control, utilizing Lipofectamine 3000 (Invitrogen). Forty-eight hours post-transfection, luciferase activity was quantified employing a dual luciferase reporter assay system (Beyotime).

### Animals

All experiments were conducted in compliance with the animal welfare guidelines sanctioned by the Animal Use and Care Committee of the Air Force Medical University. Efforts were rigorously undertaken to minimize animal distress. Male BALB/c nude mice, aged six weeks and weighing between 18-20 grams, were procured from SJA Laboratory Animal Company in Changsha, China, and were randomly allocated into various treatment groups. Prior to the commencement of the experiment, the mice were acclimated for one week in a pathogen-free animal housing facility.

### Intrathecal (i.t.) injection

We conducted intratibial (i.t.) injections following the methodology outlined in previous studies [18, 19], with minor modifications. Specifically, BALB/c nude mice (7 weeks old, 6 per group) were anesthetized using sodium pentobarbital, and their right legs were sterilized with an iodine solution. Lentivirus-infected MDA PCa 2b cells were drawn into a 25 μl Hamilton syringe (Alltech Associates, Switzerland) equipped with a 27-gauge needle. The needle was then inserted into the cortex of the anterior tubercle using a rotating ‘drilling’ motion, penetrating 3-5 mm beyond the bony cortex [18]. Subsequently, 25 μl of the cell suspension, containing 2.5 × 10^6^ cells per inoculum, was gradually injected into the bone marrow cavity. Following injection, the needle was carefully withdrawn, and bone wax was applied to seal the injection site.

### Micro-computed tomography (micro-CT) analysis

To evaluate bone mass, a micro-computed tomography (micro-CT) system (GE Healthcare, USA) was employed, following a modified version of previously established protocols [20, 21]. Mice were euthanized, and the right tibia was excised and fixed in 4% paraformaldehyde (PFA) for 24 hours. Subsequently, 7 mm specimen blocks were prepared. These specimens underwent imaging via a 120 min micro-CT scan, utilizing a resolution of 8 µm, a current of 80 µA, and a voltage of 80 kV. The acquired data were analyzed using Micview V2.1.2 software (GE Healthcare), and the mean bone mineral density (BMD) was quantified.

### In vivo imaging

Tumor progression was assessed utilizing the IVIS Spectrum system (Perkin Elmer), following the methodology outlined in a prior study [22]. Specifically, a dose of 150 mg/kg of D-luciferin, dissolved in 100 µl of PBS, was administered intraperitoneally, and the subject was anesthetized with 1.5% isoflurane before imaging. Tumor growth and burden were quantified using Living Image software (Perkin Elmer), employing bioluminescence imaging techniques.

### Orthotopic injection

Male BALB/c nude mice (7 weeks old, 5 per group) were anesthetized with sodium pentobarbital. The abdominal area was aseptically prepared for surgical intervention with an iodine solution. A surgical incision was then performed in the lower abdominal region to access the prostate, followed by injection of MDA-PCa-2b-luc cells overexpressing miRNA (1 x 10^6^ cells in 10 µL) into each dorsal prostate lobe utilizing a Hamilton syringe equipped with a 27-gauge needle.

### Prostate tissue small EVs (sEVs) isolation

The isolation of small extracellular vesicles (sEVs) from tissue was conducted following established protocols [23, 24]. Briefly, prostate tissue was sectioned and finely minced. Subsequently, the tissue underwent enzymatic digestion using collagenase D (2 mg/ml, Roche) and DNase I (40 U/ml, Roche). The mixture was then subjected to filtration and differential centrifugation to facilitate the isolation of extracellular vesicles (EVs). The centrifugation process involved sequential steps: initially, the supernatant was centrifuged at (1) 300 g for 10 minutes to remove cells, (2) 2000 g for 20 minutes to eliminate cell debris, (3) 16,500 g for 30 minutes to isolate large EVs, and finally, (3) 120,000 g for 2.5 hours to isolate sEVs.

### Droplet digital PCR (ddPCR)

To examine the expression of miRNAs within EVs, RNA was isolated utilizing the Total Exosome RNA and Protein Isolation Kit (Invitrogen). Digital droplet PCR (ddPCR) was conducted following established protocols [25, 26]. In summary, the total RNA was applied to Saphire or Opal chips, and ddPCR was executed on the Naica Crystal Digital PCR System (Stilla Technologies) employing the ddPCR Eva-Green Mix (APExBio). The resulting data were analyzed using the Crystal Reader (Stilla Technologies).

### Statistical analysis

All data are presented as means ± standard deviation. Comparisons were performed using Students *t* test and one-way ANOVA, as appropriate, using GraphPad Prism version 7.0. *P* < 0.05 was considered statistically significant (*, *P* < 0.05; **, *P* < 0.01; ***, *P* < 0.001).

## Results

### miR-375 and miR-148a-3p promote osteoblast differentiation *in vitro* and *in vivo*

MC3T3-E1 cells were cultured in medium containing 10% conditioned medium. The osteogenic induction medium served as a positive control, while the standard medium for MDA PCa 2b (F12K) was used as the control group. Compared to the F12K group, the conditioned medium derived from MDA PCa 2b cells significantly enhanced osteoblast differentiation, as evidenced by Alizarin Red S staining (Fig. 1A). This effect was attenuated by the application of the exosomse generation inhibitor GW4869, suggesting that PCa-derived extracellular vesicles (PCa EVs) partially facilitate osteoblast differentiation.

**Figure 1.**
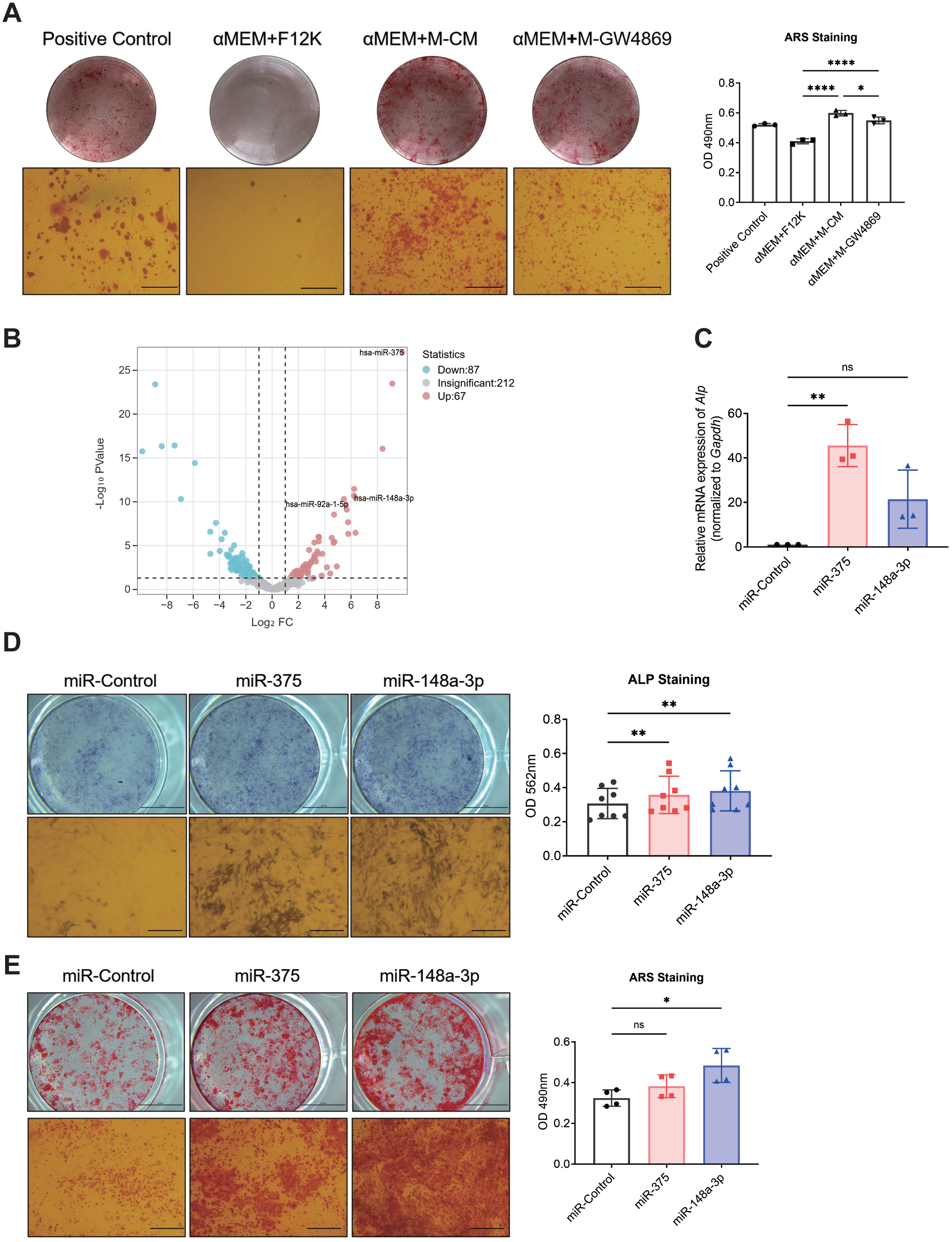
miR-375 and miR-148a-3p promote osteoblast differentiation *in vitro*. **A.** Alizarin Red S staining of MC3T3-E1 cells at Day 21 after coculture with indicated medium in osteogenic induction medium. Scale bars = 100 μm. **B.** Volcano plot showed miR-92a-1-5p, miR-375, and miR-148a-3p upregulated in MDA PCa 2b EVs. **C.** qPCR analysis of osteoblast markers *ALP* after a 6-day treatment with miRNA mimics. The medium was changed every 3 days. **D.** ALP staining of MC3T3-E1 cells at Day 7 after transfection with indicated miRNAs in osteogenic induction medium. Scale bars = 5 μm (upper) and 100 μm (lower). **E.** Alizarin Red S staining of MC3T3-E1 cells at Day 21 after transfection with indicated miRNAs in osteogenic induction medium. Scale bars = 5 μm (upper) and 100 μm (lower). Data were analyzed using one-way ANOVA. *, *P* <0.05; **, *P* < 0.01; ****, *P* < 0.0001.

Subsequent next-generation sequencing was employed to profile small RNAs from MDA PCa 2b EVs and RWPE-1 EVs. Differentially expression analysis (|log2(fold change)| > 2.5 and P < 0.05) revealed 87 downregulated and 67 upregulated miRNAs (Fig. 1B). In line with our previous work, miR-92a-1-5p, miR-375, and miR-148a-3p were upregulated in exosomes [14]. In our previous study, miR-92a-1-5p was shown to promote osteoclast differentiation, while also inhibiting osteoblast differentiation [14]. Following transfection, miRNA-375 induced upregulation of *ALP* mRNA expression was shown (Fig. 1C). ALP and Alizarin Red S staining demonstrated that mimics of miR-375 and miR-148a-3p significantly enhanced osteoblast differentiation post-transfection (Fig. 1D-E).

We then studied the impact of miR-375 and miR-148a-3p on bone homeostasis *in vivo* by creating stable MDA PCa 2b cell lines overexpressing these two miRNAs via lentivirus. MDA PCa 2b cells were injected into the right tibia of 7-week-old BALB/C nude mice (n=6 per group), followed by confirmation of successful injection using *in vivo* imaging on Day 1 (Fig. 2A). Tumor burden and bone homeostasis were assessed using *in vivo* imaging and micro-CT on Day 42. Significant morphological differences on bone mass were noted between the control and treated groups (Fig. 2B). Micro-CT also revealed significantly increased BMD in the Lv-375 and Lv-148a-3p groups (Fig. 2C).

**Figure 2.**
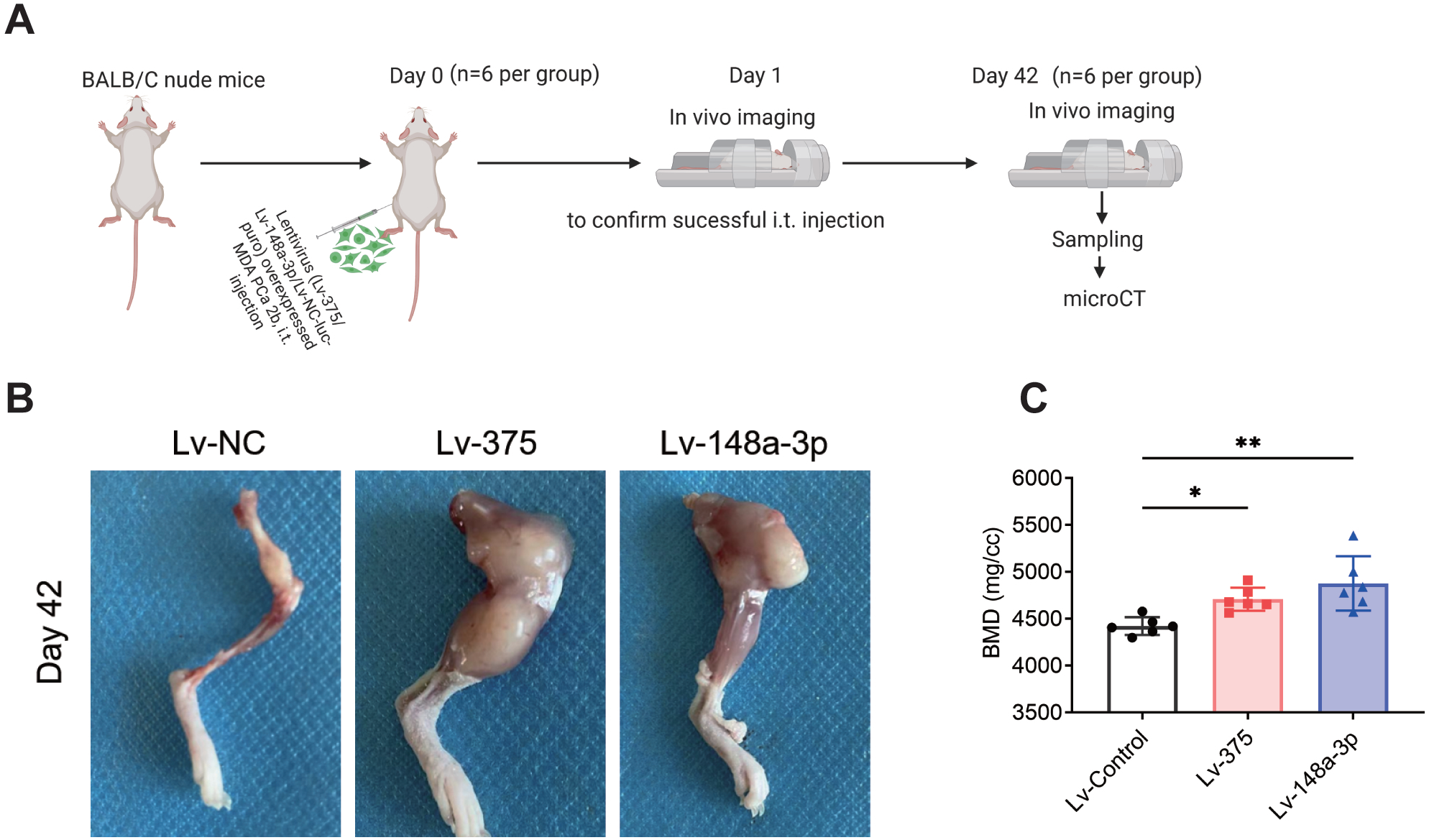
miR-375 and miR-148a-3p promote osteoblast differentiation *in vivo*. **A.** Experimental design. **B.** The representative images of bone on Day 42 after injection. **c.** Six weeks after injection of Lv-Control cells, Lv-375 cells, and Lv-148a-3p cells, micro computed tomography (micro-CT) analysis was performed to determine bone mineral density (BMD). Data were analyzed using-one way ANOVA. *, *P* <0.05; **, *P* < 0.01.

### miR-375 and miR-148a-3p promote tumor growth

After injection of the lentiviral transfected cells (Lv-375 and Lv-148a-3p) into the mouse tibia, successful injection was first verified with *in vivo* imaging on Day 1, followed by a second round of imaging on Day 42 to monitor tumor development (Fig. 3A). On Day 42, significant tumor cell proliferation was observed at the tumor sites for the Lv-375 and Lv-148a-3p groups (Fig. 3B). Bioluminescence measurements taken on Days 1 and 42 revealed a substantial increase in bioluminescence intensity within the Lv-375 and Lv-148a-3p groups (Fig. 3C). Furthermore, enhanced MDA PCa 2b cell proliferation in the Lv-375 and Lv-148a-3p groups was corroborated using the CCK8 assay (Fig. 3D).

**Figure 3.**
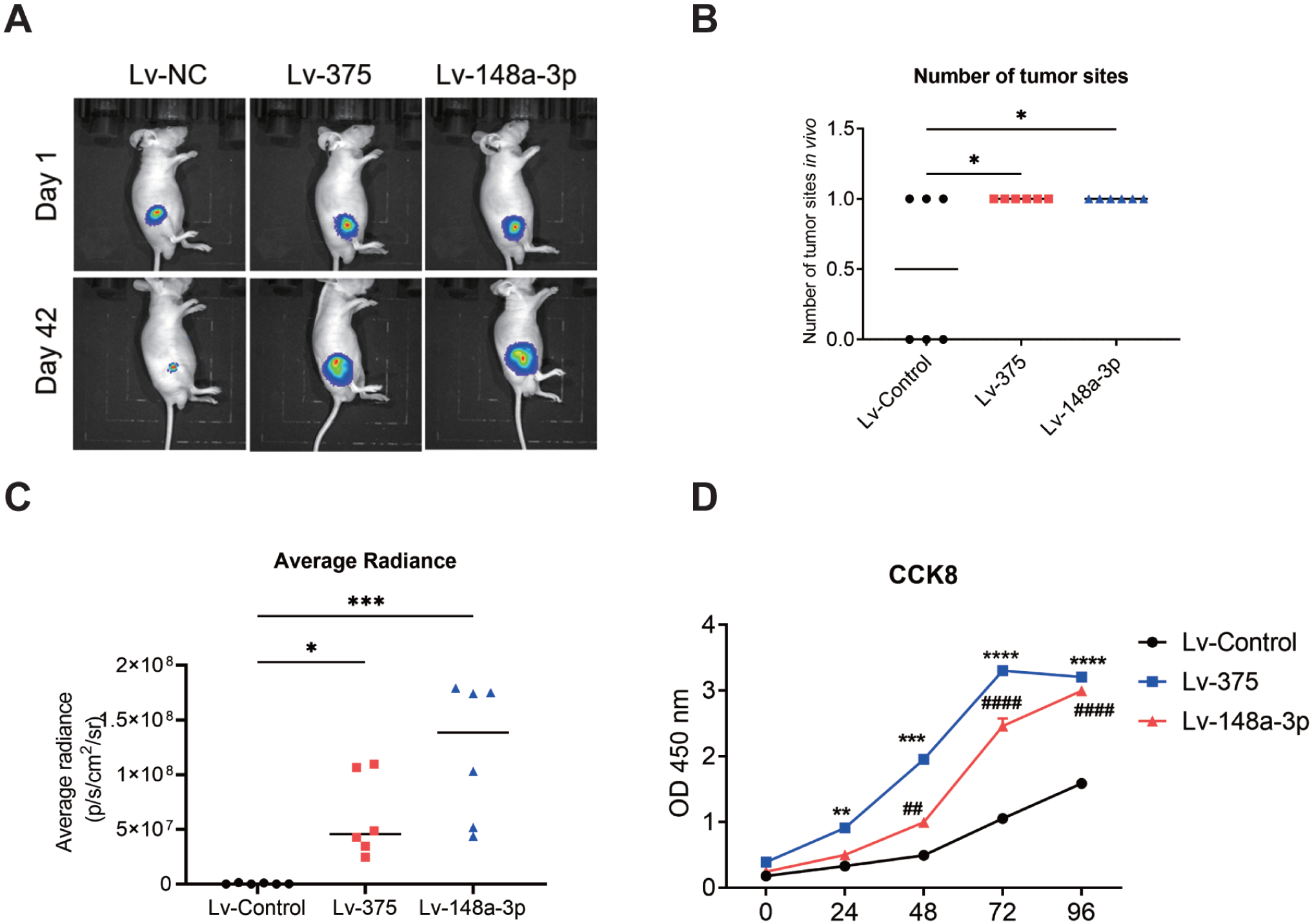
miR-375 and miR-148a-3p promote tumor growth. **A.** *In vivo* imaging of BALB/c nude mice injected with indicated cells. **B.** Day 42, a comparison of the number of tumor sites. **C.** Quantification of total tumor burden on Day 42 between the groups. **D.** CCK8 test of cell proliferation at different time points. Data were analyzed using (b-c) one-way ANOVA and (d) two-way ANOVA. *, *P* <0.05; **, *P* < 0.01; ***, *P* < 0.001 ****, *P* < 0.0001.

### miR-375 and miR-148a-3p target KLF4 to promote osteoblast differentiation and tumor growth

To elucidate the molecular mechanisms involved, we employed the online miRNA prediction software TargetScan to identify putative target genes for both miR-375 and miR-148a-3p. This analysis resulted in the identification of 251 and 749 predicted target genes for miR-375 and miR-148a-3p with an overlap of 53 genes, respectively (Fig. 4A). Among the predicted targets, the tumor suppressor transcription factor *KLF4* was identified. Subsequently, we utilized a dual-luciferase reporter assay to ascertain whether miR-375 and miR-148a-3p directly target *KLF4* (Fig. 4B). This analysis demonstrated that luciferase activity in the co-transfection group of miRNAs and *KLF4*-Wt was significantly reduced compared to the control group (P < 0.001; Fig. 4C), suggesting that *KLF4* is a direct target of miR-375 and miR-148a-3p. Forty-eight hours post-transfection with the miRNAs, *KLF4* mRNA levels were significantly reduced in MDA PCa 2b cells (Fig. 4D).Western blot analysis further revealed decreased KLF4 protein levels in MDA PCa 2b cells stably transfected with miRNA-expressing lentivirus (Fig. 4E).

**Figure 4.**
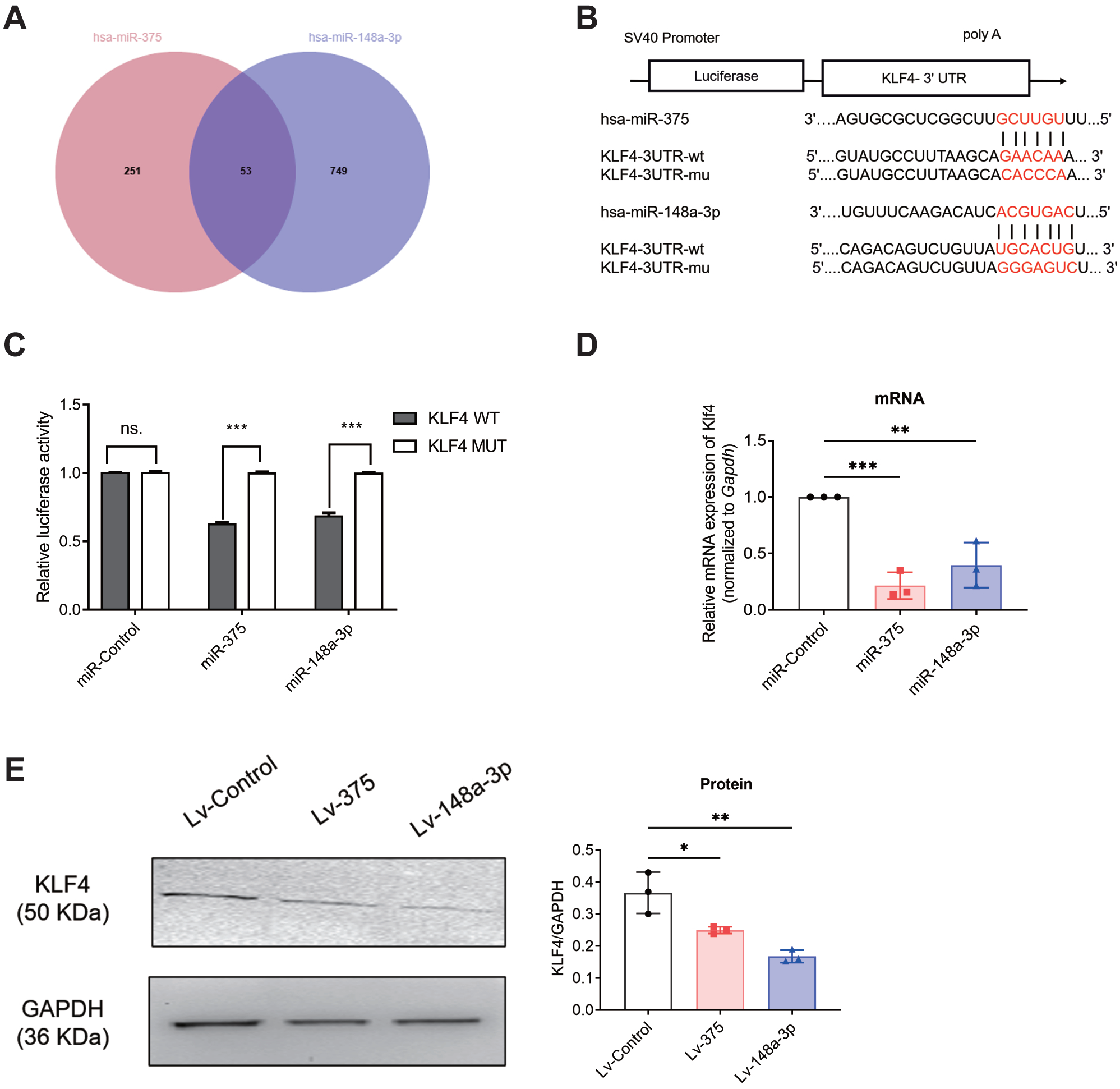
miR-375 and miR-148a-3p target *KLF4*. **A.** Venn diagram to show the potential common targets of miR-375 and miR-148a-3p. **B.** Schematic of the vectors used. **C.** Relative luciferase activity determined after 48 h of co-transfection. **D.** *KLF4* mRNA levels determined using qPCR after 48 h of transfection. **E.** KLF4 protein expression in cells stably transfected with lentivirus. GAPDH: internal control. Data were analyzed using one-way ANOVA. *, *P* <0.05; **, *P* < 0.01; ***, *P* < 0.001 ****, *P* < 0.0001.

To further elucidate the impact of *KLF4* knockdown on the differentiation of osteoblasts, we performed transfection using *KLF4* siRNA, which resulted in reduced *KLF4* expression in MC3T3-E1 cells (Fig. 5A). As expected, ALP activity and staining were significantly upregulated following the 7-day transfection period (Fig. 5B-C). Additionally, we observed an increase in MDA PCa 2b cell proliferation subsequent to KLF4 downregulation, as measured by the CCK8 assay (Fig. 5D).

**Figure 5.**
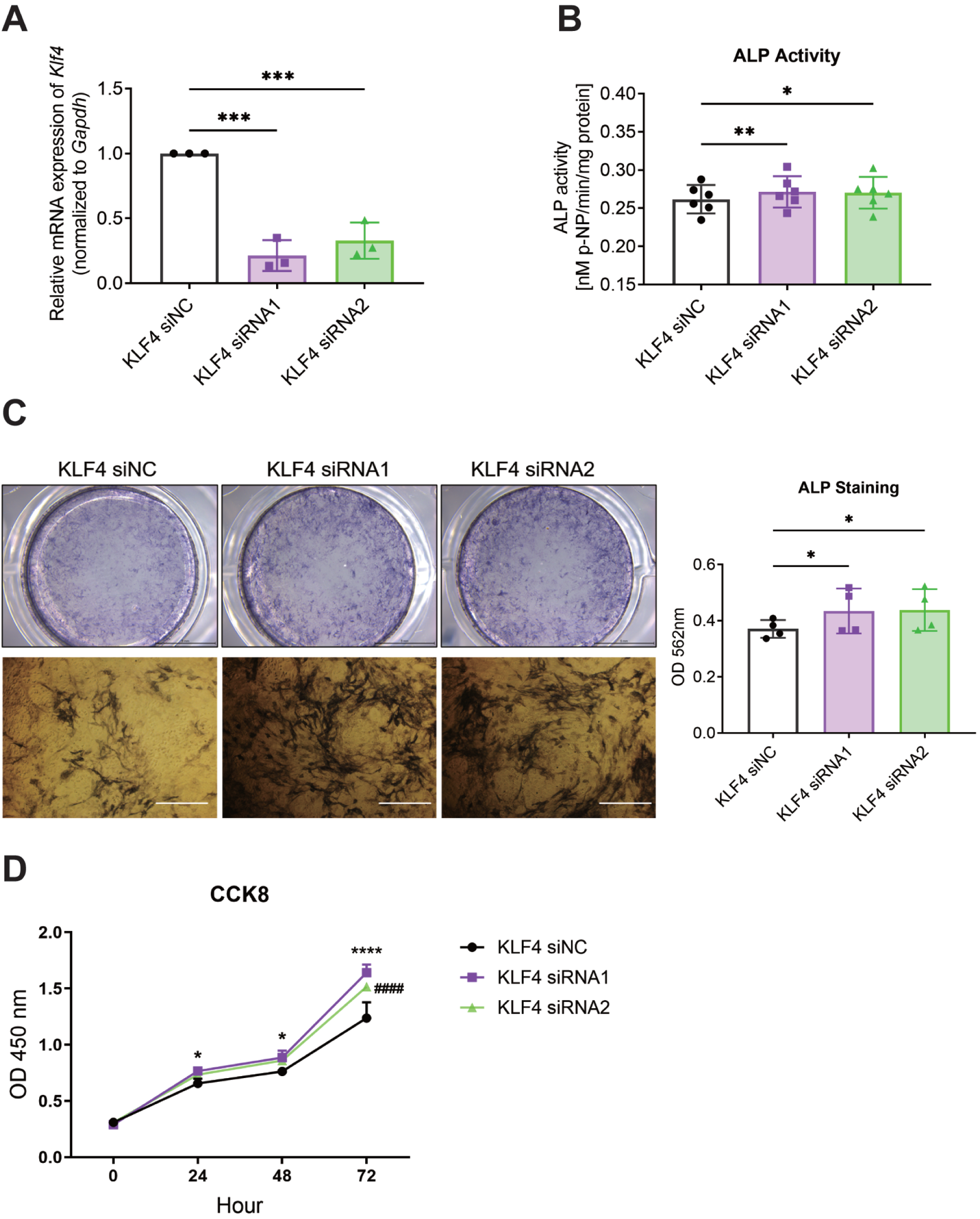
Effects of *KLF4* knockdown on osteoblast differentiation and tumor growth. **A.** qPCR analysis of mRNA expression of *klf4* in MC3T3-E1 cells transfected with indicated siRNAs and siRNA Control. **B.** ALP activity of MC3T3-E1 cells 7 days after transfection. Increased ALP staining is shown. **C.** ALP staining of MC3T3-E1 cells 7 days after transfection. Increased ALP staining is shown. **D.** CCK8 test of MDA PCa 2b cells after transfection. Increased cell proliferation is shown. Data were analyzed using (a-d) one-way ANOVA and (e) two-way ANOVA. *, *P* <0.05; **, *P* < 0.01; ***, *P* < 0.001 ****, *P* < 0.0001.

### Osteogenic and osteoclastic miRNAs may function at different time phases of bone metastasis

These findings led us to explore the coexistence of osteoblastic and osteoclastic miRNA in PCa EVs and their impact on bone homeostasis. We mimicked tumor bone metastasis by injecting miRNA-overexpressing MDA PCa 2b cells into the prostate of BALB/c nude mice (7 weeks old, n = 5 per group; Fig. 6A). After 21 and 42 days, prostate sEVs were isolated and analyzed, showing typical vesicle morphology and size (Fig. 6B-C). ddPCR revealed that miR-92a-1-5p was dominant in the early phase (Day 21), while miR-375 and miR-148a-3p were prevalent in the late phase (Day 42; Fig. 6D-G).

**Figure 6.**
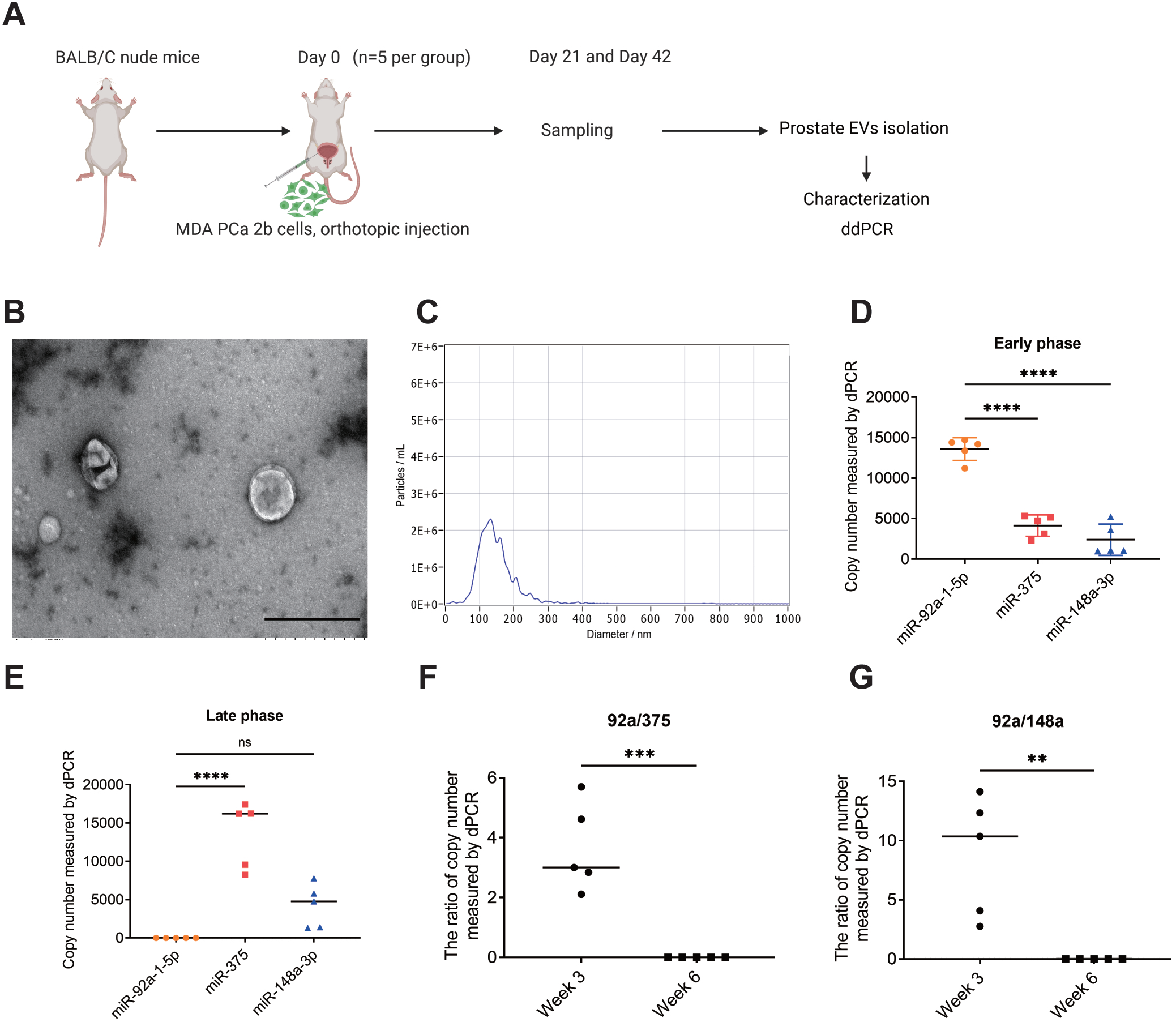
Osteogenic and osteoclastic miRNAs may regulate the PCa bone metastatic process. **A.** Experimental design. **B.** Representative transmission electron microscope (TEM) image of sEVs derived from prostate. Scale bar = 500 nm; **C.** Representative nanoparticle tracking analysis (NTA) showing average size distribution of prostate sEVs. **D.** Comparison of the expression of three miRNAs in the early phase (Day 21) of bone metastasis. **E.** Comparison of the expression of three miRNAs in the late phase (Day 42) of bone metastasis. **F-G.** Comparison of osteoclastic miRNA/osteoblastic miRNA at different time phases. Data were analyzed using (d-e) one-way ANOVA and (f-g) t-test. *, *P* <0.05; **, *P* < 0.01; ***, *P* < 0.001 ****, *P* < 0.0001.

## Discussion

In our previous work, we demonstrated that PCa EVs facilitate osteoclast differentiation while inhibiting osteoblast differentiation, with these effects being partially mediated by miR-92a-1-5p. In the present study, we found that miR-375 and miR-148a-3p, conveyed by PCa EVs, enhance osteoblastogenesis and promote tumor growth within bone tissue. Initially, we observed that the PCa-conditioned medium stimulates osteoblast differentiation, an effect partially attributable to PCa EVs. Both miR-375 and miR-148a-3p, delivered by PCa EVs, were found to promote osteogenesis *in vitro* and *in vivo*. Notably, *in vivo* imaging data further revealed that these miRNAs augment tumor growth, a finding corroborated by the *in vitro* CCK8 assay. Mechanistically, miR-375 and miR-148a-3p interact directly with *KLF4* mRNA. Additionally, we replicated the effects of miR-375 and miR-148a-3p by transfecting cells with *KLF4* siRNA, highlighting the involvement of this molecule in miR-375 and miR-148a-3p-mediated osteoblast differentiation. Finally, our findings indicate that osteoblastic and osteoclastic miRNAs coexist and function at distinct stages of PCa bone metastasis.

Since 2007, numerous studies have highlighted the pivotal role of EVs in mediating intercellular and interorgan communication [27], primarily through the delivery of their cargoes, which include proteins, lipids, DNA, mRNA, and miRNA [28-31]. Among these, miRNAs have been the most extensively studied due to their small size and high conservation across species, rendering them promising candidates for diagnostic markers and therapeutic agents in the context of EVs [32, 33]. Studies have shown that during the process of PCa bone metastasis, several miRNAs transported by PCa-derived EVs influence bone homeostasis and tumor progression. Clinical imaging data indicate that PCa bone metastasis predominantly exhibits osteoblastic characteristics [3].

Interestingly, our findings reveal that PCa EVs facilitate osteoclastic differentiation while inhibiting osteoblastic differentiation, a process partially attributed to the transfer of miR-92a-1-5p. In this study, we demonstrated that MDA PCa 2b EVs convey osteoclastic miR-92a-1-5p and osteoblastic miR-375 and miR-148a-3p at distinct stages of bone metastasis. These insights contribute to a more comprehensive understanding of the role of EV-associated miRNAs in PCa bone metastasis.

Previous studies have identified *KLF4* as an anti-tumor transcription factor that inhibits various molecular processes involved in tumorigenesis, such as cell cycle inhibition [34], tumor growth arrest [35], suppression of tumor proliferation [36], and regulation of prostate stem cell homeostasis [37]. Additionally, *KLF4* has been implicated as a regulatory factor in the processes of osteolysis and osteogenesis within bone-associated prostate tumors [38]. In the present study, we demonstrated that *KLF4* serves as a co-target gene for miR-375 and miR-148a-3p. Furthermore, the downregulation of *KLF4* via siRNAs was found to replicate the effects of miR-375 and miR-148a-3p on osteoblast differentiation and tumor growth. Previous studies have also shown that KLF4 directly interacts with the C-terminal transactivation domain of β-catenin, thereby mediating the inhibition of Wnt/β-catenin signaling [39]. This suggests that miR-375 and miR-148a-3p from PCa EVs might promote osteoblast differentiation and tumor growth via the KLF4-Wnt/β-catenin pathway, warranting further study.

We explored why osteoblastic and osteoclastic miRNAs coexist in PCa EVs. Small RNA sequencing confirmed the presence of osteoblastic miRNA-375/miRNA-148a-3p and osteoclastic miRNA-92a-1-5p in these EVs. Using a bone metastasis model, we injected PCa cells into the prostate and isolated sEVs, detecting miRNA expression with ddPCR. We observed higher levels of osteoclastic miR-92a-1-5p during the early stages of bone metastasis, and osteoblastic miR-375 and miR-148a-3p during the later stages. This suggests that microRNAs delivered by extracellular vesicles in PCa are involved in bone metastasis, initially promoting osteoclastic activity and subsequently facilitating osteoblastic activity. Consequently, these microRNAs contribute to creating space for the tumor in the early stages and support tumor growth in the later stages. Further research is required to elucidate these mechanisms.

The metastasis of tumors to the bone is a complex, multi-step process. The maintenance of bone homeostasis, mediated by the interactions between osteoblasts and osteoclasts, is essential for the bone microenvironment. Recent perspectives suggest that the tumor microenvironment within the bone acts as the “soil” facilitating tumor metastasis, while the primary tumor, or “seed,” can influence the tumor microenvironment from a distance. The progression of tumor growth in bone during the later stages of metastasis further amplifies this feedback loop. In this study, we concentrated on PCa small extracellular vesicle-delivered microRNAs. Notably, we discovered that different miRNAs within the same subpopulation of extracellular vesicles have distinct roles in bone homeostasis and the metastatic process.

## Conclusion

In conclusion, this study clarified that miR-375 and miR-148a-3p delivered by PCa EVs promote osteoblast differentiation and tumor growth via targeting *KLF4*. We also initially demonstrated that miRNAs delivered by PCa EVs participate in the bone homeostasis and bone metastasis process.

## List of abbreviations

ddPCR: droplet digital
PCR: EVs: extracellular vesicles
FBS: fetal bovine serum
miRNAs: microRNAs
MOI: multiplicity of infection
NC: Negative Control
NTA: nanoparticle tracking analysis
PBS: Phosphate-buffered saline
PCa: prostate cancer
sEVs: small EVs
TEM: Transmission electron microscopy.

## DECLARATIONS

### Ethics approval and consent to participate

The research was approved by the Animal Ethics Committee at Air Force Medical University, China.

### Consent for publication

The author approved the manuscript for publication.

### Data availability

All the original data are available upon reasonable request from correspondence author.

### Authors’ contributions

L.Y. designed the experiments, performed the experiments, analyzed the data, wrote the draft manuscript and provided funds.

## Acknowledgements

I acknowledge Prof. Xiaoke Hao at Air Force Medical University, Prof. Bingdong Sui at Air Force Medical University, Prof. Lei Zheng at Southern Medical University, Prof. Roger Olofsson Bagge at Gothenburg University, Prof. Toshima Parris at Gothenburg University, and Dr. Ting Ding at Xi’an Jiaotong University for their support and scientific advices for this work.

